# SynSeg: Generating Synthetic Datasets for Accurate Subcellular Segmentation with U-net

**DOI:** 10.1101/2025.02.07.637194

**Authors:** Zhengyang Guo, Kaiming Xu, Zi Wang, Jingyi Ke, Jingwen Huang, Yuqi Ye, Guangshuo Ou

## Abstract

Accurate segmentation of subcellular components is crucial for understanding cellular processes, but traditional methods struggle with noise and complex structures. Convolutional neural networks improve accuracy but require large, time-consuming, and biased manually annotated datasets. Here, we developed SynSeg, a pipeline that generates synthetic training data to train a U-net model for subcellular structure segmentation, eliminating the need for manual annotation. SynSeg leverages synthetic datasets with variations in intensity, morphology, and signal distribution to deliver context-aware segmentations, even in challenging imaging conditions. We demonstrate SynSeg’s superior performance in segmenting vesicles and cytoskeletal filaments from culture cells and live *C. elegans*, outperforming traditional methods such as Otsu’s thresholding, ILEE, and FilamentSensor 2.0. Additionally, SynSeg effectively quantified disease-associated microtubule morphology in live cells, uncovering structural defects caused by mutant Tau proteins linked to neurodegenerative diseases. These results highlight the potential of synthetic data-driven approaches to advance biological segmentation and enhance microscopy techniques.

**Significance Statement:** This study introduces a novel approach for accurately segmenting cellular structures, such as microtubules and vesicles, using synthetic datasets and advanced deep learning techniques. By leveraging a U-Net model trained on thousands of artificially generated images, our method eliminates the need for labor-intensive experimental data and simplifies the data creation process. Importantly, it incorporates noise and variability into the training datasets to make the model more robust and biologically relevant.

Our findings demonstrate that the model can successfully identify cellular components, paving the way for its application in real-world microscopy images. This innovation has the potential to accelerate discoveries in cell biology by providing an efficient, scalable tool for analyzing complex cellular structures, even in challenging imaging conditions.

## Introduction

Accurate segmentation of subcellular components is crucial for advancing our understanding of intracellular transport, structural organization, and cell signaling ^1–3^. However, the complexity and variability of biological images present significant challenges in achieving both precision and generalizability in segmentation methods. Traditional unsupervised techniques, such as thresholding (e.g., Otsu’s)^4^ and edge-enhancement methods like ILEE^5^, are often applied to simpler tasks. While computationally efficient and label-free, these approaches struggle in noisy environments, under uneven lighting, and with complex shapes, as they lack the contextual awareness needed to capture subtle structural details.

In contrast, machine learning methods, particularly convolutional neural networks (CNNs), have revolutionized image segmentation by learning complex patterns from labeled data ^6–8^. While CNNs deliver high accuracy, they are constrained by the need for large, manually annotated datasets, which are time-consuming, labor-intensive, and susceptible to bias ^9, 10^. Simulation-based datasets offer an alternative by generating synthetic images through physical modeling ^10, 11^, thus eliminating manual annotation. However, these datasets require significant prior expertise, complex modeling pipelines, and considerable computational resources, limiting their scalability and reproducibility.

To address these limitations, we developed SynSeg, a pipeline that automatically generates synthetic datasets to train a U-net architecture^7^ for subcellular structure segmentation. U-net is a deep learning model designed for image segmentation, which includes an encoder that extracts features and a decoder that reconstructs the image, highlighting key structures. U-net’s “skip connections” preserve fine details while integrating higher-level features, making it highly effective for segmenting complex biological images. This architecture excels in noisy or irregular images and can learn directly from data without the need for manual feature extraction, making it a powerful tool for biological studies.

By training a basic U-net on 1,000 synthetic images, we achieved strong performance in segmenting the cytoskeleton and vesicles—without requiring fine-tuning on microscopy data. With its flexibility, ease of use, and robust performance, SynSeg offers a scalable, efficient solution for biological image segmentation, demonstrating how synthetic datasets can significantly reduce the labor and complexity involved in developing high-performing models.

## Results

### Stepwise Tuning of SynSeg for Vesicle Segmentation

Vesicle segmentation presents significant challenges due to the variation in vesicle size and fluorescence intensity, especially for vesicles below the optical resolution limit^9^. To address this, we developed a synthetic image generation pipeline for training vesicle segmentation models (Fig. 1 a). The pipeline begins by creating a background that consists of random fluorescence and Gaussian noise. Vesicles are simulated by placing dots of varying radii and opacities onto the background. Each dot is then subjected to a Gaussian blur filter, with a randomly selected sigma value, to mimic defocus effects (Fig. 1 a). Ground truth masks are generated by labeling each vesicle during the creation process.

**Figure 1:**
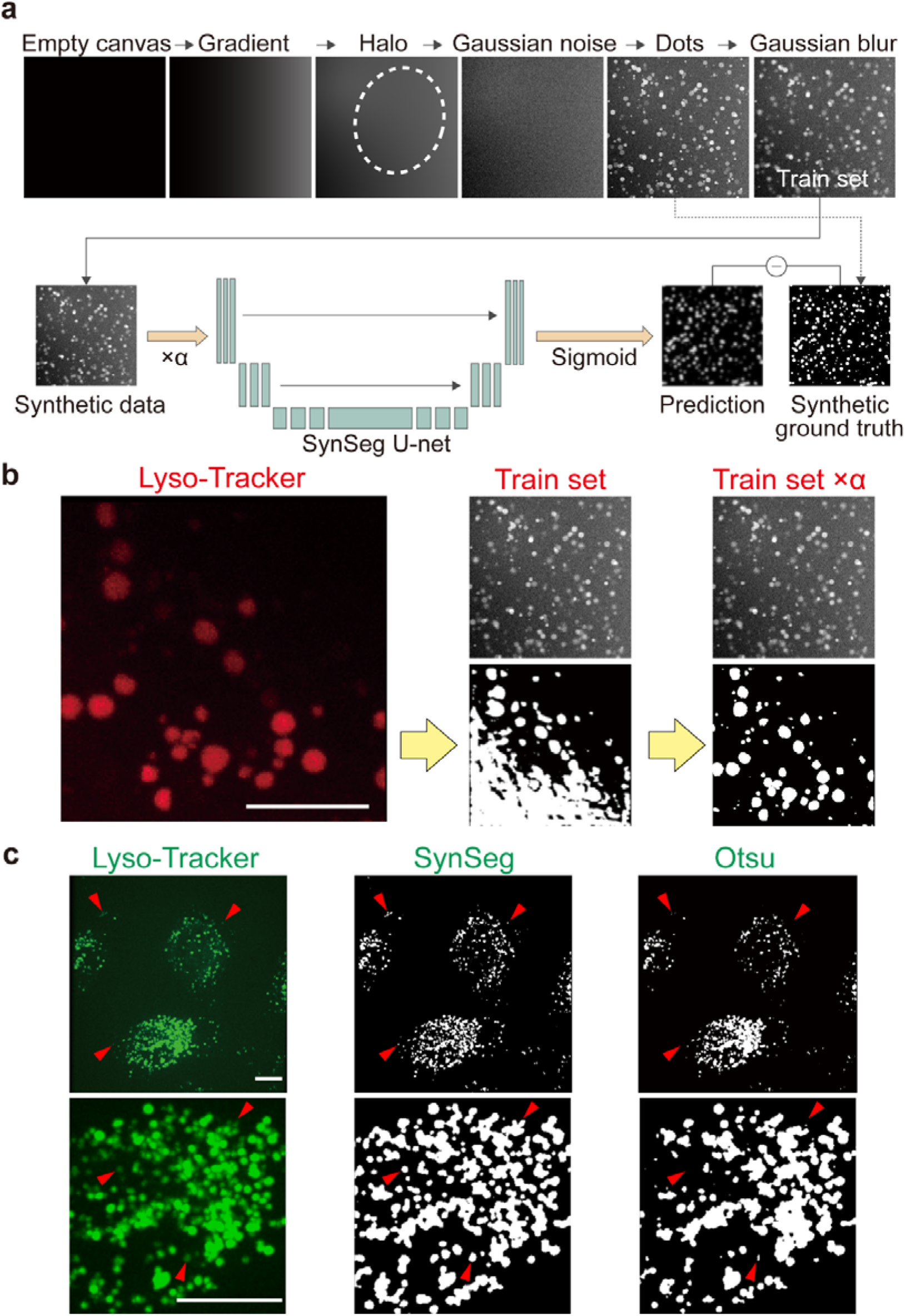
Stepwise tuning of Vesicle segmentation model. (a) Synthetic vesicle dataset generation and model architecture. The training set was generated on a 512×512 black canvas. First, a gradient with a random direction was applied, followed by a random halo to simulate the variability of background fluorescence. Gaussian noise was then added to the background, and white dots with varying opacity and radii were overlaid to simulate vesicles. Each dot was processed with a Gaussian blur filter to simulate defocus. A U-net model was trained using this synthetic vesicle dataset and corresponding ground truth masks. (b) Vesicle segmentation in *C. elegans* lysosome-related organelles. Spinning-disk confocal imaging of *C. elegans* lysosome-related organelles stained with Lyso-Tracker. The background fluorescence interfered with the prediction of vesicle masks. To improve model performance, a random scale factor (α) was applied to the pixel values as a data augmentation technique, which enhanced vesicle segmentation accuracy. (c) Vesicle segmentation in HeLa cells. Spinning-disk confocal imaging of HeLa cells stained with Lyso-Tracker (Green). The bottom row is an enlarged portion of the image above it. Arrowheads indicate vesicles missed by Otsu’s thresholding but successfully captured by the SynSeg model. Scale bars: 10 μm.

After synthesizing the vesicle dataset, we trained a basic U-net model. Initially, the model struggled to distinguish small vesicles from the background fluorescence, indicating overfitting to the synthetic dataset. To mitigate this issue, we applied a data augmentation technique by randomly scaling the pixel values within the range of α = [0.25, 1.0] each time a training image was loaded. This intensity scaling adjustment encouraged the model to rely more on morphological features and signal distribution rather than intensity alone. As a result, the model successfully improved its segmentation of small vesicles, including those that were obscured by background fluorescence (Fig. 1 b).

Next, we compared SynSeg to a widely used unsupervised thresholding method, Otsu’s thresholding ^4^, using HeLa cells stained with Lyso-Tracker (Green). While Otsu’s thresholding captured most vesicle details, SynSeg outperformed it by leveraging prior knowledge embedded in the synthetic training set. SynSeg was able to accurately identify dim and small vesicles while preserving morphology and intricate details, even in regions of high fluorescence intensity (Fig. 1 c, d).

### Stepwise Tuning of SynSeg for Cytoskeleton Segmentation

Building on the success of vesicle segmentation, we extended our approach to cytoskeleton segmentation, a particularly challenging task due to the complex network of filaments, variations in filament thickness, differences in fluorescence intensity, and the presence of subpixel-thin filaments. A new synthetic dataset was created to represent cytoskeleton features (Fig. 2 a). The generation process begins by drawing a polygon to simulate the cell shape, followed by the addition of Bézier curves with three random control points to represent filaments. Random black and white dots are added to simulate cellular textures, and a local blur is applied to the entire cell region. To replicate the thin filaments observed in cytoskeleton imaging, images are initially created at a high resolution of 2048×2048 and then downsampled to 1024×1024. This ensures that some filaments are thinner than one pixel and have non-integer positions, which helps the model learn to segment thin filaments at a subpixel scale. Ground truth masks are generated by labeling each filament during the creation process.

**Figure 2:**
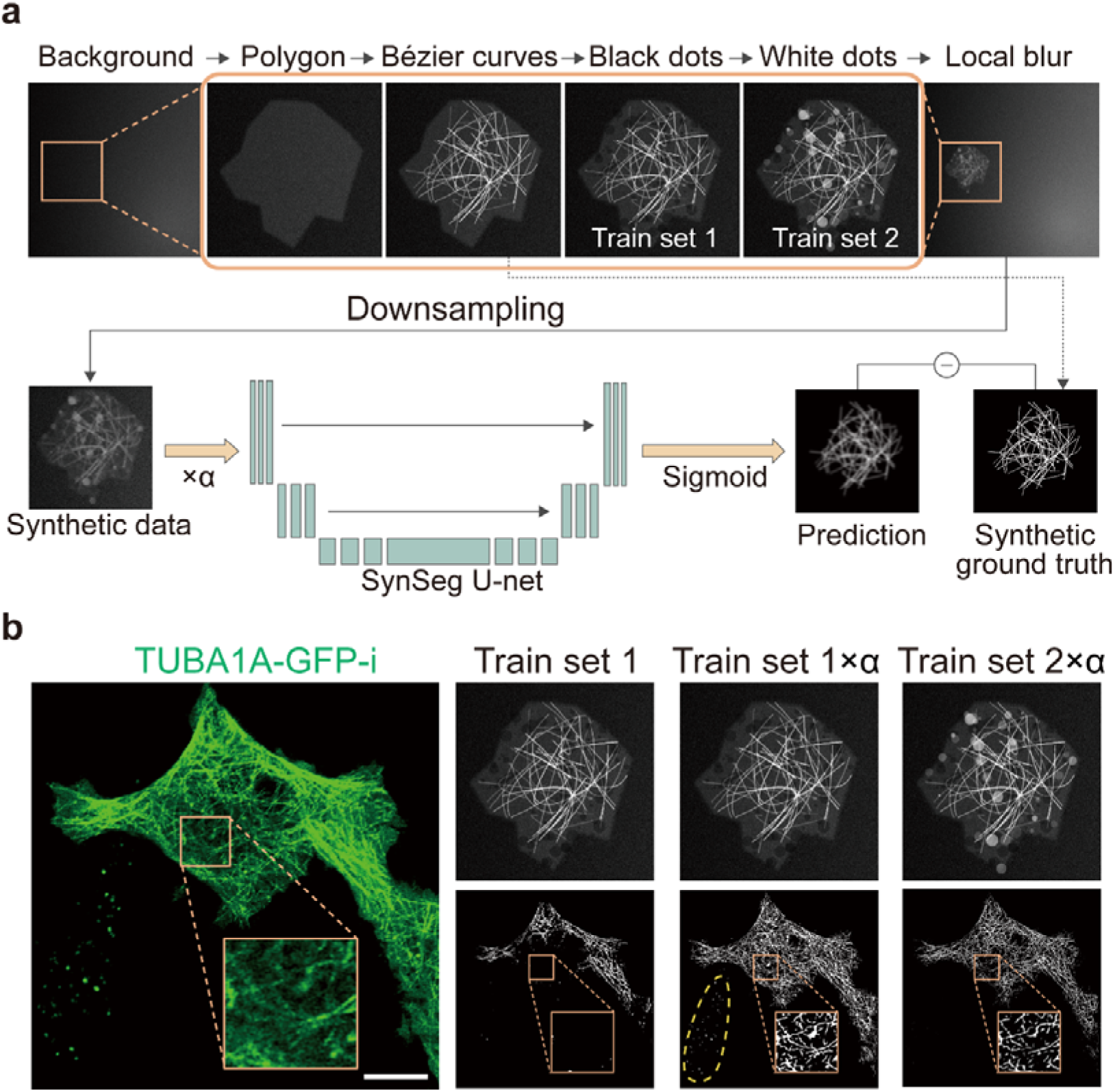
Stepwise tuning of Cytoskeleton segmentation model. (a) Synthetic cytoskeleton dataset generation and model architecture. The synthetic training set was generated on a 2048×2048 black canvas. Background generation followed a similar approach to the vesicle dataset. A random-shaped polygon was drawn on the background, within which Bézier curves were placed to simulate filaments. Black dots were randomly added to the polygon area to create Train Set 1. Subsequently, additional white dots were used to create Train Set 2. A U-net model was trained using this synthetic cytoskeleton dataset downsampled to 1024×1024, with corresponding ground truth masks for both train sets. (b) Cytoskeleton segmentation on Airyscan-based imaging of TUBA1A labeled with split-GFP. Train sets 1 and 2 were generated as described in (a). The left panel shows the original image. The top panel of each column displays the corresponding train set, with the predicted mask result shown at the bottom panel. Dashed lines highlight textures and dots unrelated to microtubule signals. Scale bars: 10 μm.

We initially trained the model on the cytoskeleton dataset without augmentation, but it failed to detect weak and thin filaments, which were not captured in the segmentation results (Fig. 2 b). To address this, we applied the same data augmentation technique used in vesicle segmentation—scaling the pixel values with a random factor α. This adjustment allowed the model to detect weak and thin filaments, including those with subpixel-level thickness. However, the model also began capturing unrelated noise, such as random textures, which impacted the segmentation quality. (Fig. 2 b) To overcome this, we introduced random white dots to the dataset. These dots helped the model distinguish filament-like signals from other unrelated shapes in the image. With this additional training data, the model was able to successfully segment the cytoskeleton, accurately identifying thin filaments while minimizing noise interference.

### Comparison with Widely-Used Segmentation Methods

We compared SynSeg with widely-used methods for cytoskeleton segmentation, including: (1) automated image-processing techniques such as Otsu’s thresholding ^4^, and (2) semi-automated tools like ILEE ^5^ and FilamentSensor 2.0 ^12^. Given that both ILEE and FilamentSensor 2.0 require significant parameter tuning, we used ILEE with default settings and optimized FilamentSensor 2.0 for best performance. ILEE and FilamentSensor 2.0 include their own preprocessing steps, while Otsu’s thresholding uses the raw image as input. For SynSeg, the image is either resized linearly or padded to match the input shape of the neural network, with no additional preprocessing applied.

We tested SynSeg, Otsu’s thresholding, ILEE, and FilamentSensor 2.0 on Airyscan-based imaging of TUBA1A-labeled HeLa cells using a split-GFP system ^13^. SynSeg successfully captured nearly all filament structures, distinguishing filament signals from noise and reconstructing complex networks. In contrast, FilamentSensor 2.0 struggled with thin filaments and condensed bundles, while ILEE performed slightly better than Otsu’s thresholding but still failed to generate accurate masks. Otsu’s thresholding was hindered by unbalanced fluorescence intensity, especially in areas with high signal (Fig. 3a, b). Additionally, due to the U-net model’s ability to recognize shape and pattern, SynSeg effectively filtered out noise from fluorescent dots in the original image.

**Figure 3:**
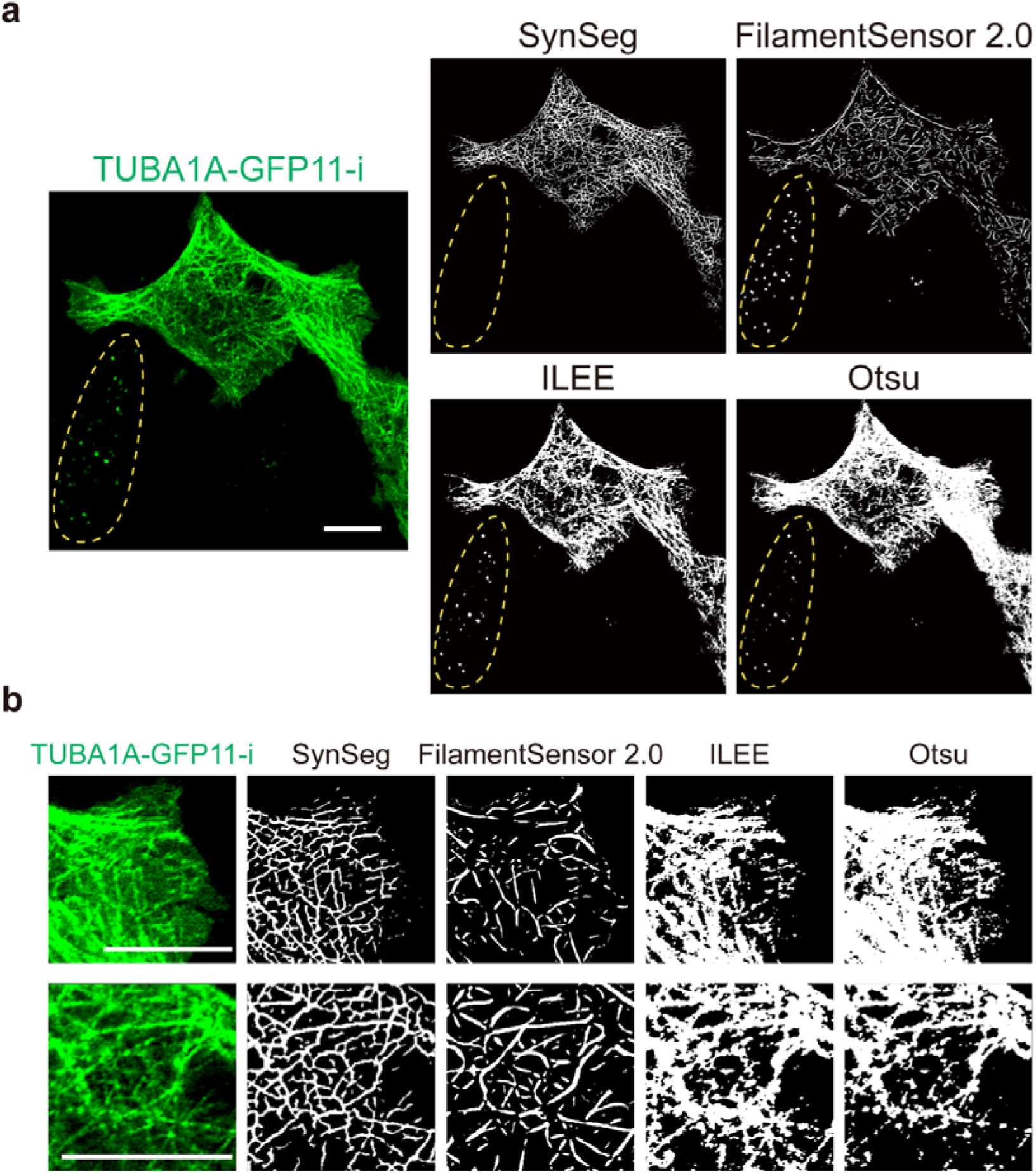
Comparative analysis of vesicle and cytoskeleton segmentation using SynSeg and other methods. (a) Cytoskeleton segmentation comparison among SynSeg, FilamentSensor 2.0, ILEE, and Otsu’s thresholding. The image was captured using an Airyscan microscope, with the microtubule network in a HeLa cell labeled by a split-GFP system. The dashed line highlights a fluorescent signal pattern that does not resemble microtubules, which was successfully filtered out by SynSeg. (b) Enlarged view of two selected regions from panel a, illustrating the performance of each method in detail. Scale bars: 10 μm.

We then tested this method on Airyscan-based imaging of split-GFP labeled beta-actin ACTB in HeLa cells^24^ (Fig. 4). In this actin filament segmentation task, SynSeg consistently outperformed the other methods. Despite the challenge of thinner filaments and greater fluorescent intensity imbalances, SynSeg accurately identified almost every filament. FilamentSensor 2.0 generated more noisy fragments compared to our method and neglected some thin filaments in its binary mask. ILEE performed better than in the previous task but still failed to generate continuous filament structures, while Otsu’s thresholding continued to struggle with high-intensity imbalances (Fig. 4a, b).

**Figure 4:**
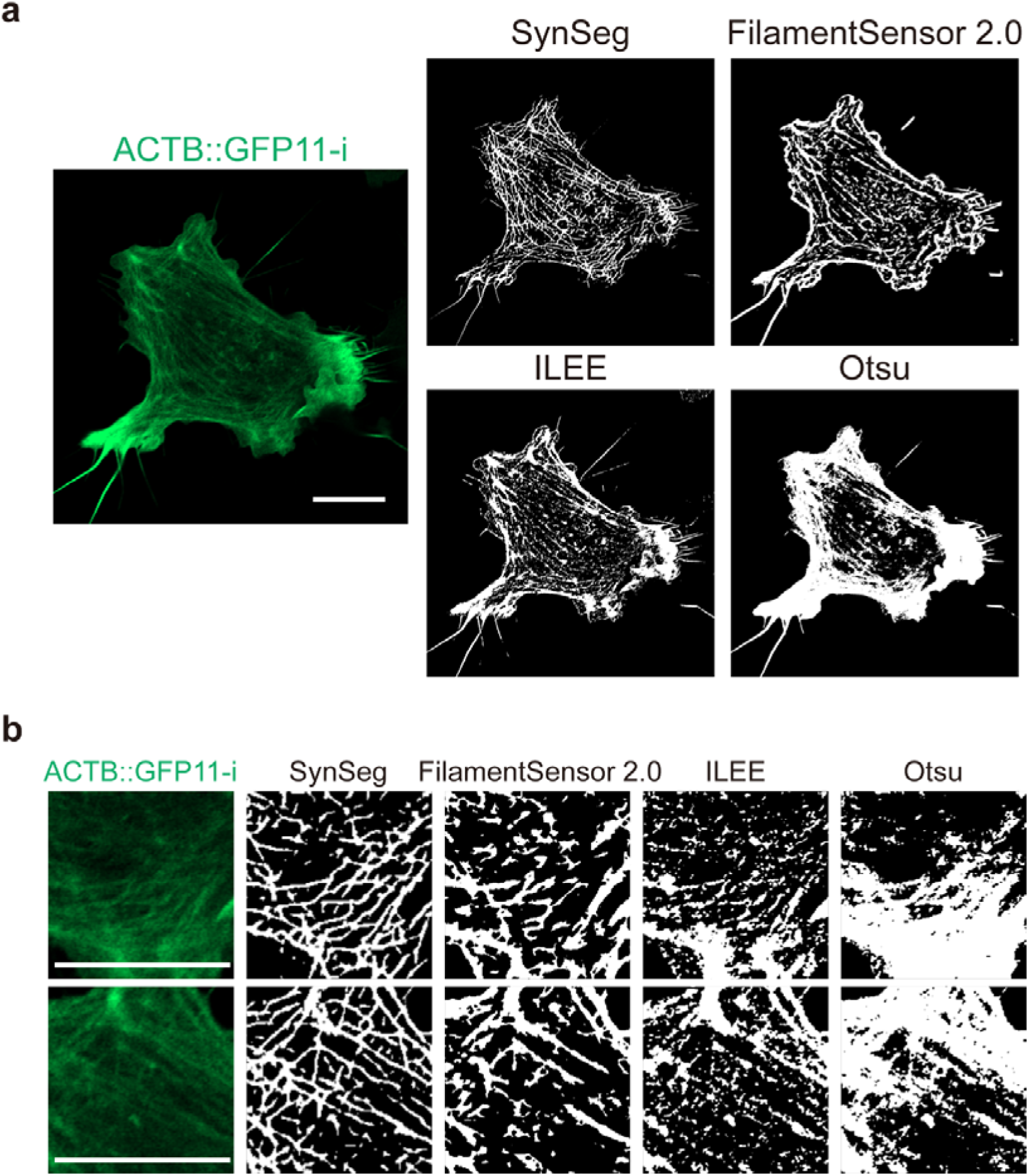
Comparative analysis of actin filament segmentation using SynSeg and other methods. (a) Cytoskeleton segmentation comparison among SynSeg, FilamentSensor 2.0, ILEE, and Otsu’s thresholding. The image was captured using an Airyscan microscope, with the action network in a B16-H10 cell labeled by a split-GFP system. (b) Enlarged view of a selected region from panel a, illustrating the performance of each method in detail. Scale bars: 10 μm.

### Cytoskeletal Dynamics in Live *C. elegans*

To evaluate SynSeg’s performance in dynamic imaging of the cytoskeleton in live animals, we applied it to spinning-disk confocal movies of epidermal microtubules in *Caenorhabditis elegans* (*C. elegans*), labeled with a *col-19* promoter-driven TBB-2-GFP11-i ^13^. These movies posed specific challenges for our model. The spinning-disk confocal system, operating at a 200 ms interval, introduced significant noise from the EM-CCD camera and provided lower resolution compared to Airyscan-based confocal microscopy. In the analysis of individual frames, FilamentSensor 2.0 failed to reconstruct detailed networks, while ILEE managed fluorescent intensity imbalances better but lacked the necessary precision. Otsu’s thresholding was once again unable to capture details in high-fluorescent intensity areas. In contrast, SynSeg successfully generated intricate masks, accurately identifying crossing microtubules (Fig. 5 a).

**Figure 5:**
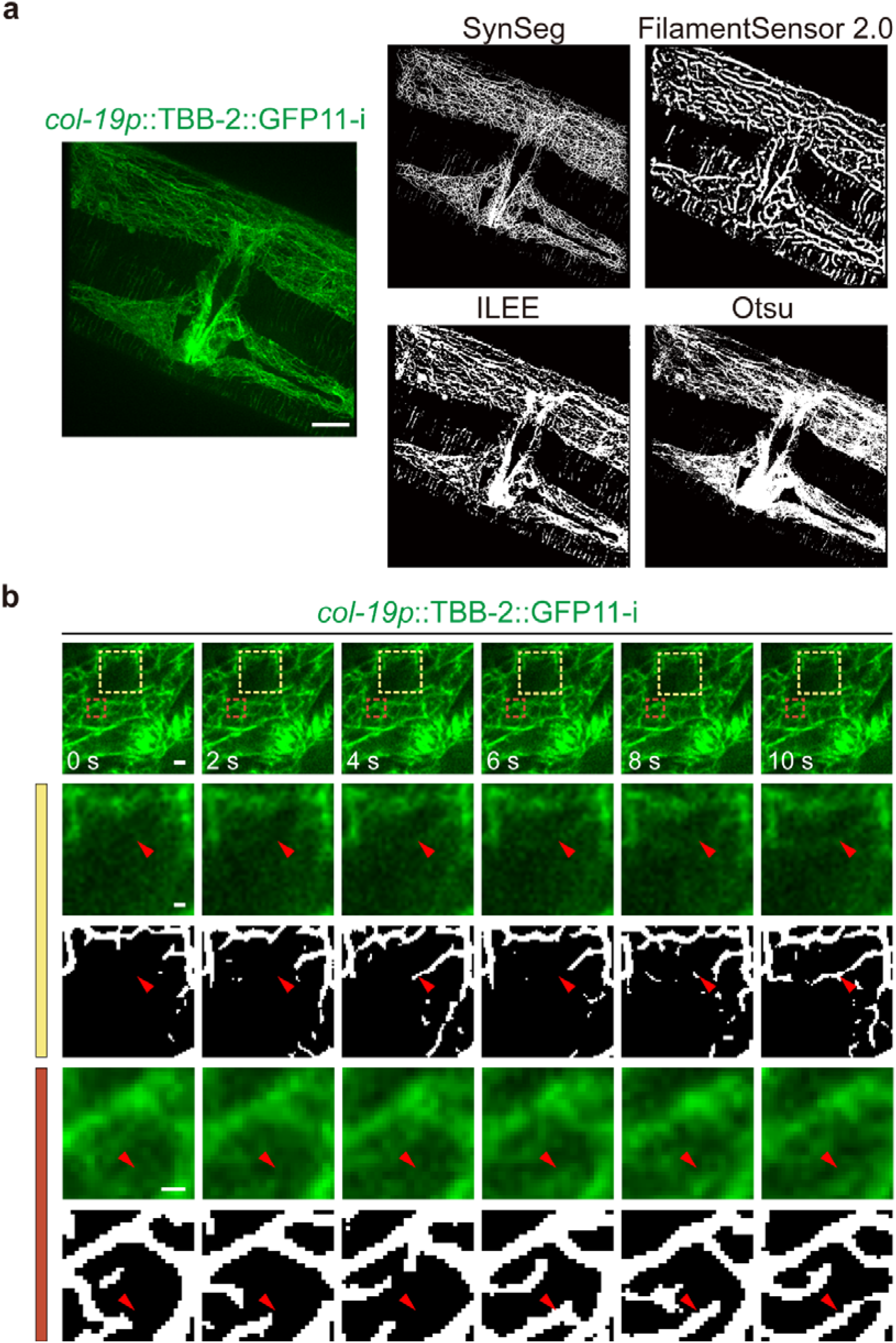
In-vivo cytoskeleton dynamic visualization. (a) *C. elegans* epidermal microtubules labeled using an internal split-GFP-tagged TBB-2 construct driven by the epidermal collagen promoter col-19p. The segmentation results from SynSeg, FilamentSensor 2.0, ILEE, and Otsu’s thresholding are shown for comparison. Images were acquired using a spinning-disk confocal microscope. Scale bar: 10 μm. (b) Analysis of a spinning-disk confocal movie depicting the dynamics of *C. elegans* epidermal cytoskeletons labeled as (a). The movie was captured at a 200 ms frame interval, with snapshots taken at time points 0, 2, 4, 6, 8, and 10 seconds. Enlarged views of areas within the yellow and brown rectangles are provided below. In the yellow and brown enlarged regions, arrowheads indicate growing fibers, potentially representing the polymerization of newly formed microtubules, which were accurately identified by the SynSeg-generated masks. Scale bars: 1 μm.

For analysis of a full movie stack, SynSeg consistently captured the dynamic movement of epidermal microtubules (Movie S1, Fig. 5 b). Notably, SynSeg was able to resolve subtle microtubule polymerization events that were imperceptible to the human eye, revealing detailed patterns of microtubule growth (Fig. 5 b).

### Quantification of disease-related microtubule-associated protein

Tau is a microtubule-associated protein essential for stabilizing microtubules and regulating their dynamics ^14^. Tau-F refers to a specific isoform of Tau protein with disease-relevant properties, often studied for its interactions with the cytoskeleton ^15^. The R406W mutant is a pathogenic variant of Tau associated with neurodegenerative diseases such as frontotemporal dementia and Alzheimer’s disease, characterized by its altered binding to microtubules and propensity to form aggregates ^16, 17^. We quantified the local fluorescent intensity of wildtype Tau-F and the R406W mutant in Airyscan-based imaging of HeLa cells using SynSeg. The accurate masks generated by SynSeg facilitated the quantification of local signals along the microtubule, minimizing the influence of cytosolic signals. Our analysis revealed distinct differences in microtubule morphology: wildtype Tau-F produced evenly distributed microtubule arrays, while Tau-F (R406W) led to a pronounced bundled structural pattern. Quantitative fluorescence analysis of local fluorescent intensity confirmed significantly higher intensity for Tau-F (R406W), indicating ectopic microtubulin bundling (Fig. 6 a, b). These results suggest that the method is effective for analyzing intensity distributions, especially in noisy or heterogeneous backgrounds.

**Figure 6:**
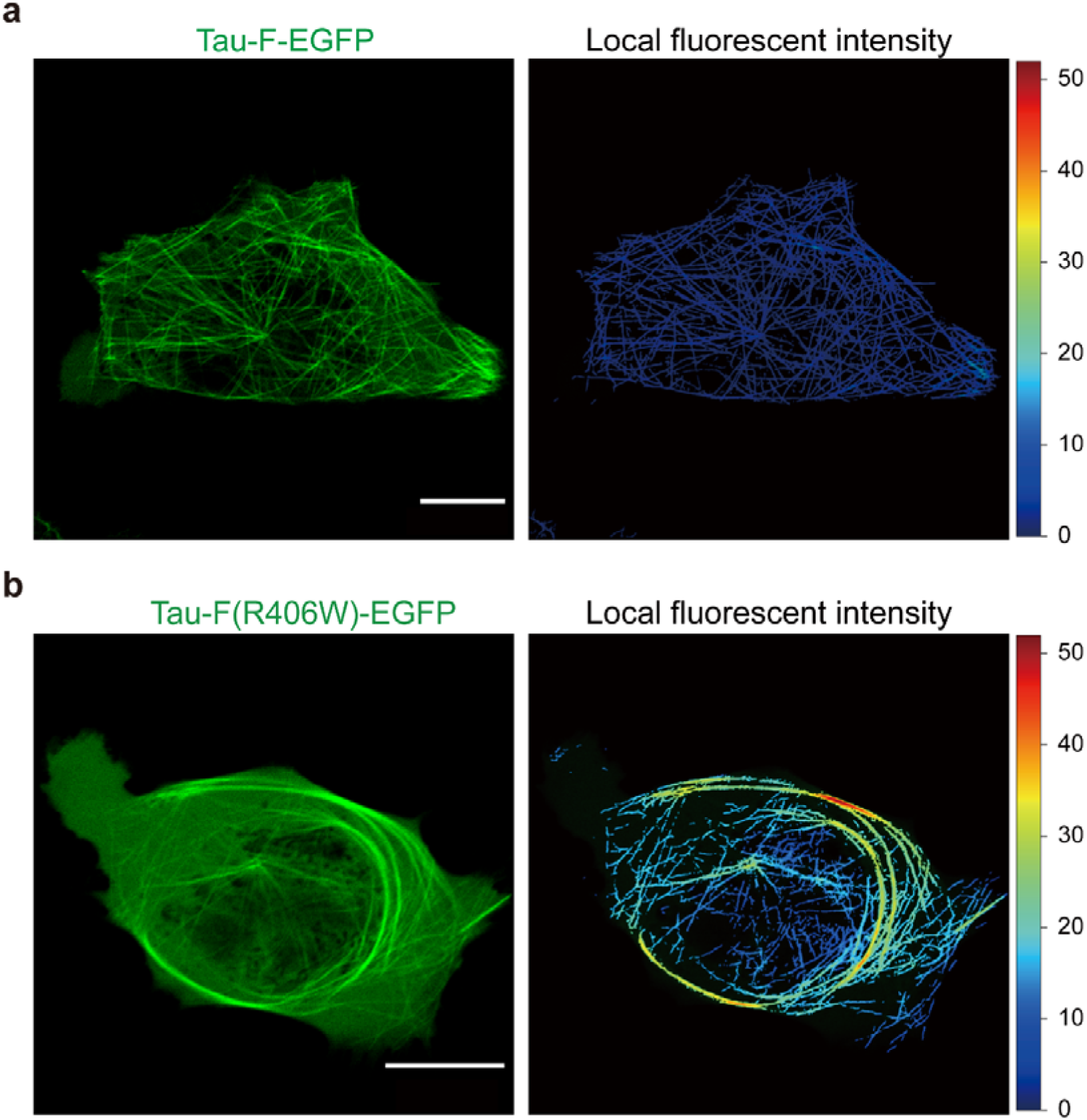
Quantification of local intensity of Tau-F and Tau-F R406W mutation. (a) Transient transfection of Tau-F-EGFP in a HeLa cell. The image was captured by Airyscan confocal. A mask was generated based on the fluorescent signal, enabling the calculation of local intensity along the microtubule fibers. (b) Transient transfection of Tau-F(R406W)-EGFP in a HeLa cell. The image was captured by Airyscan confocal. A mask was similarly generated, and the local intensity along the microtubule fibers was calculated. Scale bar: 10 μm.

Together, these results highlight the versatility and robustness of SynSeg in subcellular segmentation, from cultured cells to live animals under both physiological and pathogenic conditions. By leveraging synthetic datasets with a U-net backbone, SynSeg consistently outperformed conventional methods, excelling in scenarios with unbalanced fluorescence, thin filaments, and noisy data. This work underscores the transformative potential of synthetic datasets in advancing biological image segmentation.

## Discussion

In this study, we introduced the SynSeg approach for subcellular structure segmentation that eliminates the labor-intensive process of manual annotation while delivering robust performance. Unlike traditional methods that rely on annotated experimental datasets, SynSeg uses synthetic training data tailored to specific segmentation tasks, overcoming the limitations of physical simulation techniques, which, while providing high realism, demand detailed prior knowledge of cellular structures, microscopy techniques, and photokinetic. These methods usually involve intricate modeling workflows and substantial computational resources to capture biological complexity accurately. In contrast, SynSeg offers a simplified, adaptable solution that enables the rapid generation of biologically relevant datasets for diverse segmentation tasks.

A key strength of SynSeg is its ability to train models that generalize well to experimental data, even under challenging conditions such as unbalanced fluorescence intensities, complex geometries, and noisy imaging. By introducing random variations in intensity, morphology, and signal distribution, SynSeg enhances the model’s focus on pixel intensities, morphological features, and contextual information, leading to highly accurate and context-aware segmentation. This approach demonstrates the transformative potential of synthetic datasets in biological imaging, offering a scalable and efficient solution for complex biological studies. Furthermore, the dataset generation strategy is versatile and can be adapted for use with other neural network architectures, such as YOLO^18^ and R-CNN^19, 20^ for object detection and segmentation, or tracing algorithms like DeepSORT^21^, expanding its applicability to a wide range of tasks.

The robust performance of SynSeg highlights the power of training on fully synthetic datasets, suggesting that this approach could be extended to advanced microscopy techniques ^22, 23^. Given its strong ability to detect microtubule filaments—features that are often difficult to discern with the human eye—SynSeg demonstrates how synthetic data generation can be leveraged to train models that achieve high sensitivity to desired image patterns. While further adjustments are needed for broader applications, this method holds potential for enhancing spatial resolution and overcoming traditional imaging constraints. By harnessing fully synthetic, data-driven approaches, future models may unlock new possibilities for advancing microscopy techniques and provide unprecedented structural insights, assuming that datasets are appropriately tailored.

Our findings show that synthetic datasets, as exemplified by SynSeg, offers a transformative solution in biological imaging. The ability to design task-specific training datasets, combined with the pipeline’s simplicity and scalability, provides new opportunities to tackle longstanding challenges in cell biology and biophysics. Future directions include expanding SynSeg’s application to other subcellular structures and exploring its integration with real-time imaging platforms, particularly in scenarios where traditional segmentation methods fall short.

## Methods

### Synthetic Cytoskeleton Dataset Generation

Datasets are generated with python scripts in a 2048×2048 resolution and then down sampled linearly into a 1024×1024 picture. Here we generated 1500 pictures for training and 100 pictures for testing. A binary mask was also synthesized for ground truth to mark the position of the synthesized filaments.

### Background Generation

The background of the synthetic images was initialized with a horizontal gradient to simulate a baseline fluorescence signal (Fig. 1 a). To incorporate variability reflective of experimental conditions, a halo artifact with a randomly sampled central coordinate and intensity was overlaid onto the gradient background. This halo was designed to mimic localized fluorescence signals that might arise from uneven illumination or autofluorescence.

To further emulate real-world signal-to-noise ratio (SNR) characteristics, random noise sampled from a normal distribution was added to the image (Fig. 1 a). This noise introduces stochastic fluctuations in pixel intensity, capturing the inherent variability observed in fluorescence microscopy. Together, these elements create a realistic and diverse background that challenges the model to distinguish filaments from varying fluorescence conditions effectively.

### Synthesized Cell image with Filaments

A synthetic cellular area was represented by a white polygon with 8 to 15 edges, randomly generated to mimic the irregular shapes of real cells. Filaments within the cell were simulated using Bézier curves, defined by three control points located within the polygonal boundary. The Bézier curve was parameterized using the equation:

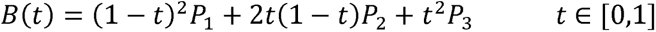

To ensure the curves appeared as continuous filaments, a dilation operation was applied to the sampled points using a 3×3 kernel. The dilation was iterated 1–2 times to introduce variability in filament thickness, mimicking the structural diversity observed in cytoskeletal networks.

Each cellular area was populated with 40 such filaments, resulting in a complex and realistic representation of cytoskeletal patterns. This procedural generation approach captures the intricate geometries of cytoskeletal filaments while allowing for controlled variability in their properties, such as thickness and orientation.

### Texture generation and blur

To replicate the appearance of vesicles and other organelles commonly observed in microscopy images, random transparent spheres with varying black and white intensities were overlaid within the synthetic cellular area. These spheres introduced additional structural complexity, mimicking the heterogeneous internal composition of cells.

To enhance realism and simulate the indistinct edges often encountered in microscopy, a Gaussian blur filter was applied to the entire cellular area. The blur softened the boundaries of each cellular component, making it more challenging to distinguish individual structures. A Gaussian kernel of size 15×15 and standard deviation σ=4 was used, effectively diffusing sharp edges while preserving the overall morphology of the filaments and organelles.

This combination of random spheres and Gaussian blurring ensured the synthetic images closely mirrored the visual characteristics of actual microscopy data, providing a robust challenge for the segmentation model.

The generated images were subsequently down sampled to a resolution of 1024×1024 pixels. This down sampling step intentionally positioned some filament curves into the sub-pixel domain, resulting in filaments with thicknesses of less than one pixel. By incorporating these ultra-thin filaments, the dataset challenged the model to identify subtle patterns and improved its ability to detect fine, faint structures in real-world scenarios.

Additionally, approximately 10% of the images in both the train and test datasets were deliberately left without any filaments. This design choice aimed to enhance the model’s capability to differentiate genuine filaments from background textures or noise. By exposing the model to a diverse range of conditions—including the absence of filaments—this approach improved the robustness and generalization of the segmentation model.

### Synthetic Vesicle Dataset Generation

Background generation was the same as cytoskeleton dataset generation. Varying radii and opacities are randomly placed on the background to simulate vesicles. Gaussian blur filters, with randomly selected sigma values, are then applied to each dot to mimic varying defocus levels.

### U-net

We employed a U-net-based architecture for cytoskeleton segmentation. The U-net design was selected for its simplicity and effectiveness in biomedical image segmentation tasks. The network comprises an encoder-decoder structure with skip connections that ensure the preservation of spatial information critical for precise segmentation.

The encoder consists of three convolutional blocks, each designed to progressively downsample the input image while extracting feature representations. Each block is composed of:

A 5×5 convolutional layer with padding to capture broader contextual information. A batch normalization layer to stabilize training.

A ReLU activation function for non-linear feature extraction.

A 3×3 convolutional layer, followed by batch normalization and ReLU, for fine-grained feature refinement.

After each convolutional block, spatial resolution is reduced using 2×2 max pooling, halving the image dimensions and doubling the number of feature channels, progressing from 64 to 256 channels.

The bottleneck layer bridges the encoder and decoder paths. It consists of a convolutional block with 512 feature channels, which captures the most abstract representations of the input data.

The decoder directly mirrors the encoder. It employs a upsampling strategy combined with convolutional operations to achieve segmentation. The upsampling is performed using transposed convolutional layers with 2×2 kernels, doubling the spatial resolution. At each stage, the upsampled feature maps are concatenated with the corresponding encoder outputs via skip connections.

The final layer is a 1×1 convolution, reducing the output to a single channel that represents the segmentation mask. A sigmoid activation function is applied to generate a pixel-wise probability map for the cytoskeletal structures.

Binary Cross Entropy (BCE) loss was utilized to quantify the discrepancy between the predicted segmentation masks and the corresponding ground truth masks. The input images, originally 8-bit or 16-bit matrices, underwent a dynamic intensity scaling process designed to enhance model robustness. Specifically, a scaling factor was randomly sampled from a uniform distribution between 0.25 and 1.0. During training, each input image was multiplied by this factor, simulating variations in fluorescence intensity commonly observed in microscopy data.

This augmentation strategy was designed to encourage the model to focus on the geometric patterns of cytoskeletal filaments rather than relying on intensity levels, which can vary across experimental conditions. By introducing variability in intensity, this approach also reduces the risk of overfitting.

During prediction, only linear resizing or padding was used for preprocessing to meet the input requirements of the neural network. The final output threshold, based on the sigmoid function (ranging from 0 to 1), was manually selected to optimize performance.

### Cell Culture and Transfection

HeLa cells were grown in DMEM (Gibco) containing 10% fetal bovine serum (Yeasen) and 1% penicillin and streptomycin (Yeasen) at 37 LJ with 5% CO2. B16-F10 cells were grown in RPMI-1640 medium (Gibco) containing 10% fetal bovine serum (Yeasen) and 1% penicillin and streptomycin (Yeasen) at 37 LJ with 5% CO2. Cells were seeded on 35 mm glass-bottom confocal dishes (CellVis) the day before transfection. Cells were transfected with tuba1a-gfp11-i (200 ng) and gfp1-10 (300 ng), or *actb*-gfp11-i (200 ng) and gfp1-10 (300 ng), or Tau-F-gfp (400 ng), respectively, using Lipofectamine^TM^ 3000 Transfection Reagent (Invitrogen) following manufacturer’s instruction. The cells were incubated with cell culture medium containing 2 μg/mL Hoechst 33342 (Yeasen) and 1x Tubulin Tracker^TM^ Deep Red (Invitrogen, #T34076) at 37 °C for 30 min before imaging. For Lyso-Tracker (Green) imaging, Cells were incubated with cell culture medium containing 500 nM Lyso-Tracker (Green) (Beyotime, #C1047S) at 37 °C for 30 min before imaging.

### Live Cell Imaging

For HeLa or B16-F10 cells, imaging was performed using Zeiss LSM900 with Airyscan2 confocal microscopy (Carl Zeiss) equipped with a 63×/1.4 objective. Images were acquired at laser wavelength of 488 nm for GFP and 405 nm for Hoechst channel. Images were taken using identical settings (Airyscan mode: Airyscan SR; Scan direction: bidirectional; Scan mode: frame). For Hela cells stained with Lyso-Tracker Green, imaging was performed using Zeiss LSM980 confocal microscope equipped with a 100×/1.46 oil objective and controlled by ZEN software (Carl Zeiss). Images were acquired at laser wavelength of 488 nm for GFP channel. To ensure consistency across imaging studies, images within the same figure panel were captured using identical exposure times and processed with the same parameters using ImageJ. Optical sections were acquired at 1 μm intervals, and z-stack images were processed using the “Max intensity” projection method in ImageJ. All images shown were adjusted linearly.

### *C. elegans* Lyso-Tracker staining

Day 1 adult *C. elegans* (approximately 30 individuals) were soaked in 100 µl of M9 buffer containing 10 µM Lyso-Tracker Red (Beyotime, #C1046) to stain the intestinal tissue. The staining procedure was conducted at 20°C in the dark for 1 hour. Following staining, the worms were transferred to NGM plates seeded with fresh OP50 bacteria and allowed to recover at 20°C in the dark for 2 hours before further analysis.

### Fluorescence Imaging of *C. elegans*

*C. elegans* hermaphrodites were anesthetized with 0.1 mmol/L levamisole in M9 buffer, mounted on 3 % agarose pads, and maintained at 20 LJ. For the nematode *C. elegans*, imaging was performed using an Axio Observer Z1 microscope (Carl Zeiss) equipped with 488 and 561 laser lines, a Yokogawa spinning disk head, an Andor iXon + EM-CCD camera, and a Zeiss 100×/1.46 objective. Images were acquired at 200 ms intervals for 100× objective. Images were taken using identical settings (EM-gain: 250, exposure time: 200 ms, 30 % of max laser).

In Lyso-Tracker imaging, animals were anesthetized using 1 μM levamisole and imaged immediately. For imaging worms stained with Lyso-Tracker Red, a Zeiss LSM980 confocal microscope equipped with a 100×/1.46 oil objective and controlled by ZEN software (Carl Zeiss) was utilized. A 561 nm laser was used for excitation. To ensure consistency across imaging studies, images within the same figure panel were captured using identical exposure times and processed with the same parameters using ImageJ. Optical sections were acquired at 0.6 μm intervals, and z-stack images were processed using the “Max intensity” projection method in ImageJ.

### Local Fluorescent Intensity Calculation

To quantify local intensity in regions of interest (ROIs), we applied a sliding window approach using a binary mask to define the ROI. For each pixel, a square window (e.g., 5×5 pixels) was centered, and the statistical measures (mean, standard deviation, minimum, or maximum) of pixel intensities within the window were computed. Only pixels within the mask were considered for calculation. The resulting local intensity values were stored in a matrix of the same size as the image. For visualization, intensity values were normalized to a 0-1 scale and overlaid as a heatmap, providing insight into spatial intensity variations within the ROI.

### Comparison Cytoskeletal Segmentation Methods

Otsu’s thresholding in ImageJ was used for comparison. For the ILEE method, we used the Python library ILEE_CSK, following the default single-image analysis tutorial, with parameters set to k1 = 2.5 and k2 = 666. For FilamentSensor 2.0, we applied the default preprocessing procedure, and manually adjusted the binary mask parameters to achieve optimal performance.

## Code Availability

All script including generate the synthetic datasets and segmentation model, including parameters, were uploaded to a github repository: https://github.com/young55775/SynSeg

## Acknowledgement

This work was supported by the National Key R&D Program of China Grants 2022YFA1302700 and 2019YFA0508401; and National Natural Science Foundation of China Grants 31991190, 92254306, 32270773, 32470730, 32070706, and 32270721; and Pillars of the Nation Funding for Life Sciences, Tsinghua University. National Natural Science Foundation of China (323B200173).

